# Comprehensive analyses of 1771 transcriptome from seven tissues enhance genetic and biological interpretations of maize complex traits

**DOI:** 10.1101/2023.08.09.552713

**Authors:** Mengyu Lei, Huan Si, Mingjia Zhu, Yu Han, Wei Liu, Yifei Dai, Yan Ji, Zhengwen Liu, Fan Hao, Ran Hao, Jiarui Zhao, Guoyou Ye, Yanjun Zan

**Affiliations:** Tobacco Research Institute, Chinese Academy of Agricultural Sciences, Qingdao, CN-266000, China; Agricultural Genomics Institute at Shenzhen, Chinese Academy of Agricultural Sciences, Shenzhen, 518120, China; College of Ecology, Lanzhou University, Lanzhou, 730000, China; Key Laboratory for Bio-Resource and Eco-Environment of Ministry of Education & Sichuan Zoige Alpine Wetland Ecosystem National Observation and Research Station, College of Life Science, Sichuan University, Chengdu, CN-610065, China; Biostatistics Department, School of Public Health, University of Michigan, Ann Arbor, Michigan, 48105, USA; College of Forestry, Northwest Agriculture and Forestry University, Yangling, CN-712100, China; College of Information Engineering, Northwest Agriculture and Forestry University, Yangling, CN-712100, China; CAAS-IRRI Joint Laboratory for Genomics-assisted Germplasm Enhancement, Agricultural Genomics Institute in Shenzhen, Chinese Academy of Agricultural Sciences, Shenzhen 518120, China

**Author notes:** Corresponding authors Guoyou Ye Yanjun Zan. These authors contributed equally.

**Keywords:** GWAS, Regulatory variants, Integrative genomics, Complex trait

## Abstract

By analyzing 1771 RNA-seq datasets from seven tissues in a maize diversity panel, we explored the landscape of multi-tissue transcriptome variation and evolution patterns of tissue-specific genes, and built a comprehensive multi-tissue gene regulation atlas to understand the genetic regulation of maize complex trait. Using transcriptome-wide association analysis, we linked tissue-specific expression variation of 45 genes to variation of 11 agronomic traits. Through integrative analyses of tissue-specific gene regulatory variation with genome-wide association studies, we detected relevant tissue types and candidate genes for a number of agronomic traits, including leaf during the day for anthesis-silking interval (*GRMZM2G093210*), leaf during the day for kernel Zeinoxanthin level (*GRMZM2G143202*), and root for ear height (*GRMZM2G700665*), highlighting the contribution from tissue-specific gene expression to variation of agronomic trait. Our findings provide novel insights into the genetic and biological mechanisms underlying complex traits in maize, and the multi-tissue regulatory atlas serves as a primary source for biological interpretation, functional validation, and genomic improvement of maize.

## Introduction

Genome-wide association studies (GWAS) have identified numerous quantitative trait locus (QTL) for crop agronomic trait (Jin et al., 2023; Li et al., 2022; Lipka et al., 2013; Liu et al., 2020; Liu et al., 2021; Owens et al., 2014; Wang et al., 2020). However, fine-mapping genes underlying those associations and dissecting the underlying biological mechanisms remains a significant challenge, partially due to a lack of knowledge in whether these variants act in transcriptional or post-transcriptional level and the functional tissue. Recently, a few studies have demonstrated the power of combining GWAS, transcriptome-wide association studies (TWAS) and expression quantitative trait loci (eQTL) analysis in functionally characterizing the association signals (Lin et al., 2022; Pang et al., 2019; Zheng et al., 2020). By leveraging population-scale transcriptome variation, many genes (Kremling et al., 2019; Lin et al., 2017) and co-expression network (Schaefer et al., 2018) were revealed. Further integrative analyses of large-scale GWAS and multi-tissue regulatory datasets in human Genotype-Tissue Expression (GTEx) project (Consortium, 2020) and in farm animal Genotype-Tissue Expression project (FarmGTEx (Liu et al., 2022; Teng et al., 2022)) have revealed the potential to discovery tissue-specific regulatory variants (eQTL) and connected trait-relevant tissues or cell types to complex trait variation, paving a promising way to obtain a comprehensive understanding on the genetic regulation of complex trait.

In crops, due to a lack of population-scale multi-tissue omics data, to the best of our knowledge, no publication has comprehensively reported the causal tissue or tissue-specific regulatory variants relevant to agronomic trait variation in maize. Such a multi-tissue regulatory road map will offer valuable information for fine-mapping causal genes/variants underlying agronomic trait (Fang et al., 2020), functional characterization of the biological mechanisms (Pan et al., 2021; Yan et al., 2020; Zhao et al., 2021) and genetic improvement programs (Clark et al., 2020) in crop species. Here, we reanalyzed a multi-tissue transcriptome dataset, including 1771 RNA-seq samples from seven tissues in a maize diversity panel with 298 inbred maize lines (Kremling et al., 2018). We constructed a multi-tissue atlas of regulatory variants, quantified the contribution of tissue-specific expression to agronomic trait variation, and connected the tissue-specific expression of 11 genes to variation of 11 agronomic traits. The results presented here, for the first time, establish connections at the RNA level between tissue and agronomic trait in crops and provide an important starting point for post-GWAS functional experiments to explore genotype-phenotype relationships in crops.

## Results

### Data summary

We downloaded 1960 publicly available RNA-Seq datasets from a maize diversity panel with 298 inbred lines, including 26 stiff stalk lines, 103 non-stiff stalk lines, 77 tropical/subtropical lines, six sweet corn lines, nine popcorn lines, and 61 mixed lines (Supplementary Table S1). After quality control, 1771 datasets from seven tissues, germinating root (RT), germinating shoot (SH), base of leaf 3 (LB), tip of leaf 3 (LT), kernel (KN), adult leaf during the day (LD), and adult leaf during the night (LN, Fig 1A) were kept for downstream analysis. Meanwhile, we downloaded 4,263,832 single-nucleotide variants (SNPs) from Kremling et al (Kremling et al., 2018) and kept 4,260,521 SNPs after quality control (Materials and Methods).

**Fig 1.**
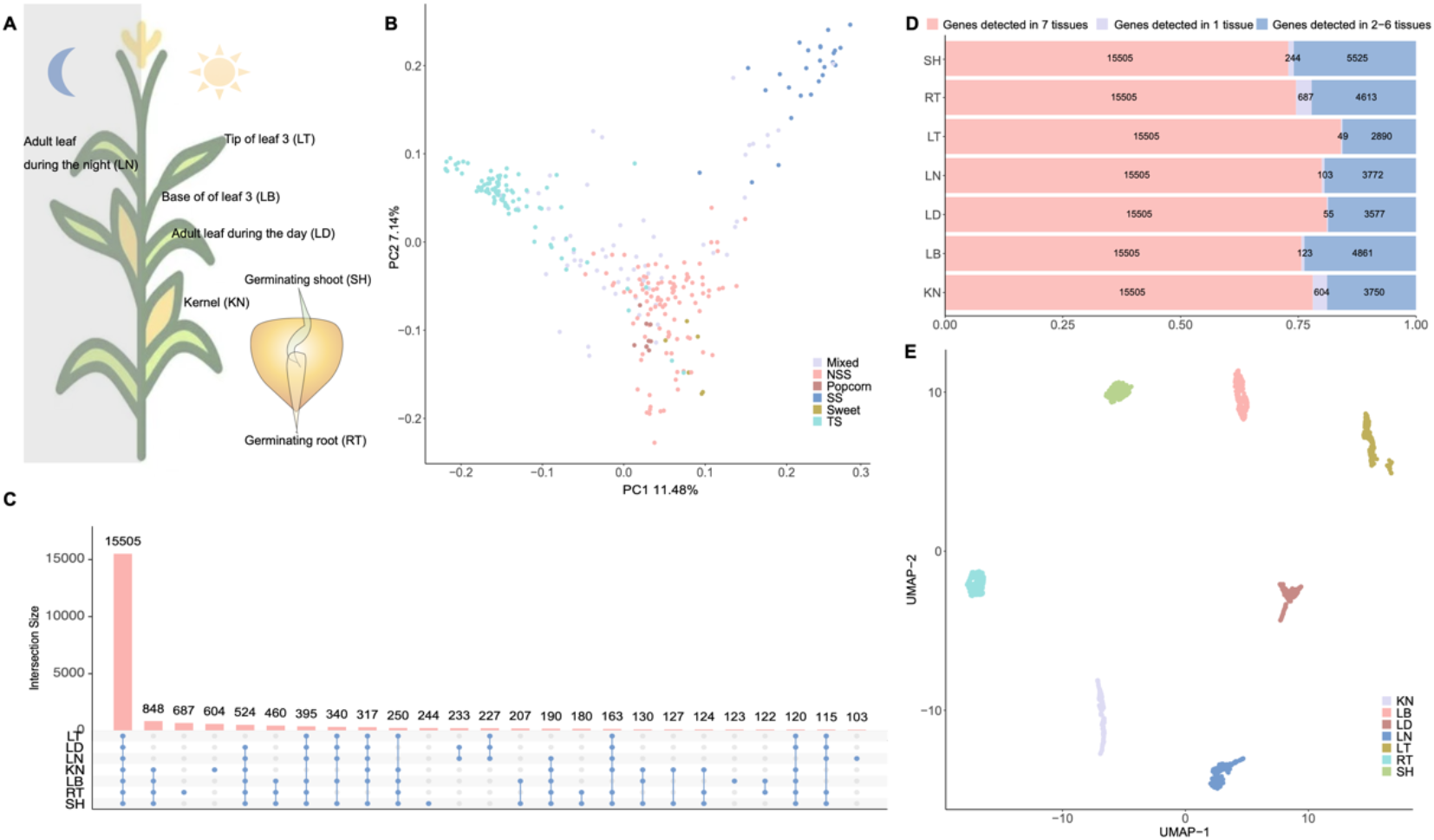
Gene expression diversity across seven tissues in the maize diversity panel. (**A**) Graphical illustration of the seven tissues analyzed in our study. (**B**) Principal component analysis of the IBS matrix estimated from 4,260,521 SNPs. (**C**) The number of core and dispensable genes in each tissue. (**D**) Overlap of genes among seven tissues. (**E**) The UMAP project of seven tissues based on 15,505 genes detected in all samples.

### Multi-tissue gene expression diversity in a maize diversity panel

The first principal component of identity by state (IBS) matrix, estimated from genome-wide SNPs, explained 11.48% (Fig 1B) of the overall variation, indicating the presence of moderate population structure. We detected an average of 19,913 genes (median = 19,859, ranging from 18,444 to 21,274 with normalized counts > 0 in more than 50% lines) across seven tissues, amount to 24,399 genes with detectable expression in seven tissues. To explore the multi-tissue gene expression diversity, we first performed all pairwise comparisons for the number of expressed genes across seven tissues. Overall, 15,505 genes (64%) were detected in all tissues (Fig 1C), while 1.31% genes were uniquely detected in one tissue (Fig 1D) on an average level. When projecting the expression matrix of 15,505 genes detected in seven tissues using Uniform Manifold Approximation and Projection (UMAP, Fig 1E), seven tissues were separated. These results indicated that tissue specificity rather than the genetic distance dominated expression variation in this dataset.

### Detection and functional characterization of the tissue–specific genes

We detected tissue-specific expressed genes one tissue at a time based on the rank of *t*-statistics and expression fold change (Materials and Methods, Supplementary Table S2). There were a number of genes (from 39 to 317, median =185) being highly expressed in each tissue across all the inbred lines, and some of them had biological functions relevant to the corresponding tissue (Fig 2A, Supplementary Fig S1). For example, *GRMZM2G045318*, a protein transport protein Sec61 beta subunit being conserved from archaea and bacteria to eukaryotic cells (Zhao and Jantti, 2009), was specifically expressed in kernel. In addition, *GRMZM2G001750*, encoding alpha/beta-Hydrolases (ABH) superfamily protein has been proved to function as bona fide hormone receptors in the strigolactone (a novel carotenoid-derived plant hormone), and gibberellin response pathways (Mindrebo et al., 2016) (Fig 2A), was specifically expressed in leaf. In addition, the number of tissue-specific expressed genes obtained at population-scale was significantly lower than that obtained from a single reference accession B73 (Supplementary Fig S1), suggesting that tissue-specific expression varies considerably among different genetic backgrounds.

**Fig 2.**
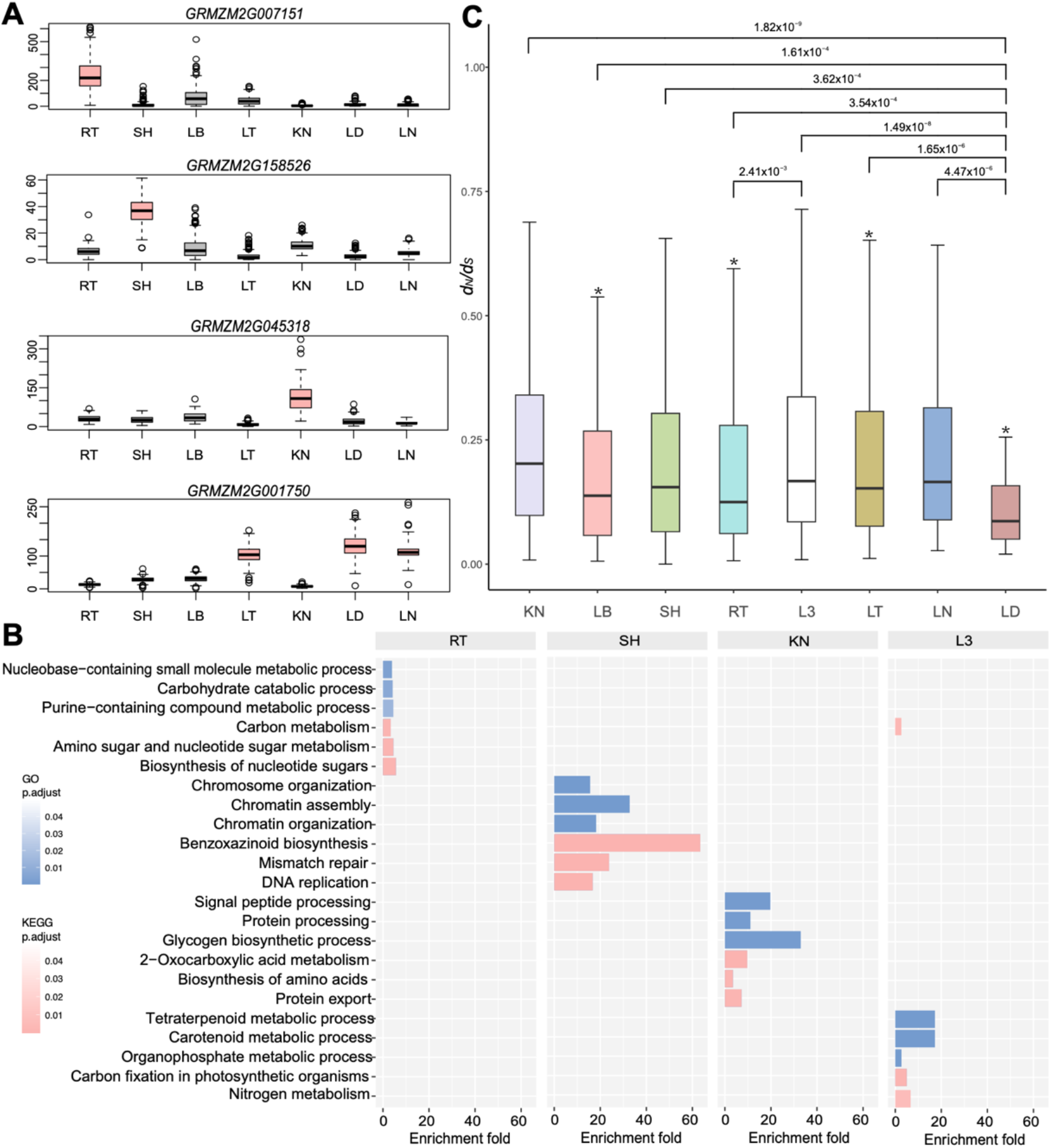
Characteristics of the tissue-specific gene in maize. (**A**) Examples of selected tissue-specific genes in RT (*GRMZM2G007151*), SH (*GRMZM2G58526*), KN (*GRMZM2G045318*), and leaf, including LT, LD, and LN (*GRMZM2G001750*). Y-axis is the gene expression. (**B**) Gene Ontology (GO) enrichment analysis and Kyoto Encyclopedia of Genes and Genomes (KEGG) of tissue-specific genes (the top 5% of genes based on *t*-statistics and median of expression). The value for each bar is the fold of enrichment. (**C**) *d_N_/d_S_* ratio between maize and teosinte for tissue-specific genes across seven tissues (L3 stands for LT, LN, and LD). *P*-values were calculated using Student’s *t*-test. For each tissue, we compared tissue-specific genes of this tissue against the remaining genes using a two-tailed *t*-test; “*” indicates *P* < 0.01

Overall, the number of tissue-specific expressed genes were functionally enriched in tissue-relevant pathways or biological processes (Fig 2B, Supplementary Fig S2, Supplementary Table S3). For example, KN-specific expressed genes were enriched for glycogen biosynthetic process (FDR adjusted *P*-value= 7.31 × 10^-4^, enrichment fold= 32.94, Fig 2B), signal peptide processing etc. (FDR adjusted *P*-value= 2.06 × 10^-4^, enrichment fold= 19.76, Fig 2B). Leaf-specific expressed genes were significantly enriched for carbon fixation in photosynthetic organisms (FDR adjusted *P*-value= 8.21 × 10^-4^, enrichment fold= 4.98 Fig 2B), nitrogen metabolism (FDR adjusted *P*-value= 6.22 × 10^-3^, enrichment fold= 6.64, Fig 2B) etc.

To explore the evolutionary conservation of tissue-specific genes, we explored *d_N_/d_S_* ratios of orthologous genes between maize and teosinte in seven tissues. LD-specific genes displayed the lowest *d_N_/d_S_* ratios (median = 0.16), while KN-specific genes had the highest *d_N_/d_S_* ratios (median = 0.20, Fig 2C). It is likely that, in constrained tissues (e.g. LD, being conserved among plants), tissue-specific genes tended to evolve slowly, whereas, in the relaxed tissues (e.g., KN, being diversified among seed plant), tissue-specific genes evolved more rapidly, revealing the importance of tissue-driven evolution.

### Genetic basis of multi-tissue gene expression diversity

To explore the genetic basis of multi-tissue gene expression diversity, we performed a comprehensive catalogue of the genetic variants underlying expression diversity in seven tissues and built a multi-tissue atlas of regulatory variants. In total, we detected 5760 eQTL for 7127 genes (eGene) in six tissues (Fig 3A, kernel was excluded due to a significantly lower number of detected eQTL, Supplementary Table S4). The number of eGenes varied significantly from one tissue to another, ranging from 1506 in LT to 2543 in RT (median = 1762.5). The majority of eGenes had one eQTL, while very few eGenes showed up to six eQTL (Fig 3B). Across all tissues, the number of *cis*-eQTL (1 Mb of the target gene’s transcription start site, TSS) was significantly larger than that for *trans*-eQTL (Fig 3C), which was in consistent with previous eQTL mapping reports (Li et al., 2013).

**Fig 3.**
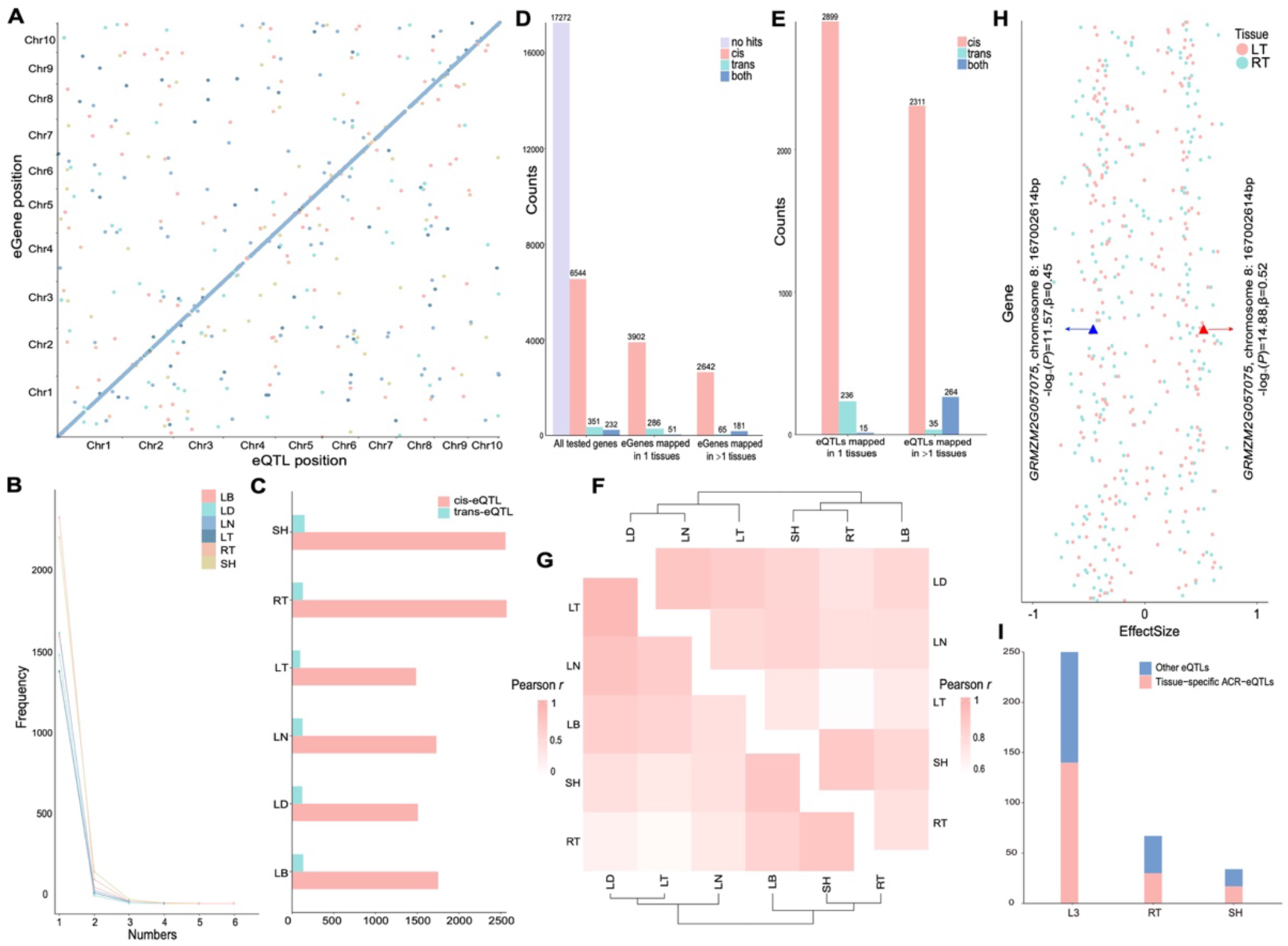
Summary of the eGWAS results. (**A**) A multi-tissue atlas of maize regulatory variants constructed from six tissues (Kernel was excluded due to a significantly lower number of detected eQTL). X-axis is the location of the eQTL, while the y-axis is the position of the associated gene. (**B**) The number of eQTL associated with one gene in each of the six tissues. (**C**) The number of *cis* and *trans* eQTL identified in each tissue. (**D**) Tissue specificity of eGenes. (**E**) The number of eQTL mapped in one and more tissues. Illustration of the pairwise Pearson correlation estimated from gene expression (**F**) and eQTL effect sizes (**G**). Tissues are grouped by hierarchical clustering. (**H**) Estimated eQTL effect sizes (x-axis) in RT and LT for all the eGenes (y-axis). Triangle symbols represent the eQTL for *GRMZM2G057075*.

To explore the similarities and differences in gene expression regulation across tissues, we compared the overlap and effect sizes of detected eQTL. 59% (4239/7127) eGenes showed eQTL in only one tissue (Fig 3D), and 55% (3150/5760) eQTL were detected in one tissue (Fig 3E), suggesting that gene expression regulation was generally tissue-specific. This was supported by the high correlation (Regression R^2^=0.58) between tissue similarity estimated from expression matrix (Fig 3F) and effect sizes (Fig 3G). In addition, the sign and direction of eQTL effect sizes changed considerably across tissues. For example, between the two most distinctive tissues, RT and LT (Pearson’s *r*=0.09, Fig 3H), 19.7% (45/228) eQTL showed effect sizes in opposite directions. Although, a single eQTL at chromosome 8: 167,002,614bp for *GRMZM2G057075* (homologous to *Arabidopsis thaliana* abiotic stress response repressor CLB (de Silva et al., 2011)) was simultaneously detected in RT and LT, the G allele increased CLB expression in RT but decreased that in LT.

### Multi-tissue transcriptome-wide association analysis highlighted the relationship between tissue-specific gene expression and variation of agronomic trait

To estimate the contribution of multi-tissue transcriptome variation to agronomic trait variation, we calculated the proportion of phenotypic variance explained (PVE) by the 24399 transcripts using a linear mixed model, one tissue at a time (Fig 4A, Materials and Methods). In this analysis, 11 agronomic traits, measured on the same population (Peiffer et al., 2014), were analyzed, including growing degree-days to silking (DTS_GDD_), growing degree-days to anthesis (DTA_GDD_), growing degree-days to anthesis-silking interval (ASI_GDD_), days to silking (DTS), days to anthesis (DTA), anthesis-silking interval (ASI), plant height (PH), ear height (EH), distance between PH and EH (PE), ratio between EH and PH (EP), and ratio between PH and days to anthesis (PD). Overall, there was considerable variation in the amount of variance explained by each tissue (median from 36.74% to 61.55%, Fig 4A). Taking DTS as an example, the SH transcriptome explained 75.45% of the phenotypic variation, while the LN transcriptome only explained 34.33%. Across six tissues (ranging from 29.73% to 52.88%), the SH transcriptome explained the highest phenotypic variance (median=61.55% for the 11 agronomic trait), followed by LB transcriptome (median=53.99%). These results revealed a variable contribution from tissue-specific transcriptome diversity to agronomic trait variation and highlighted the importance of SH transcriptome variation in agronomic trait variation.

**Fig 4.**
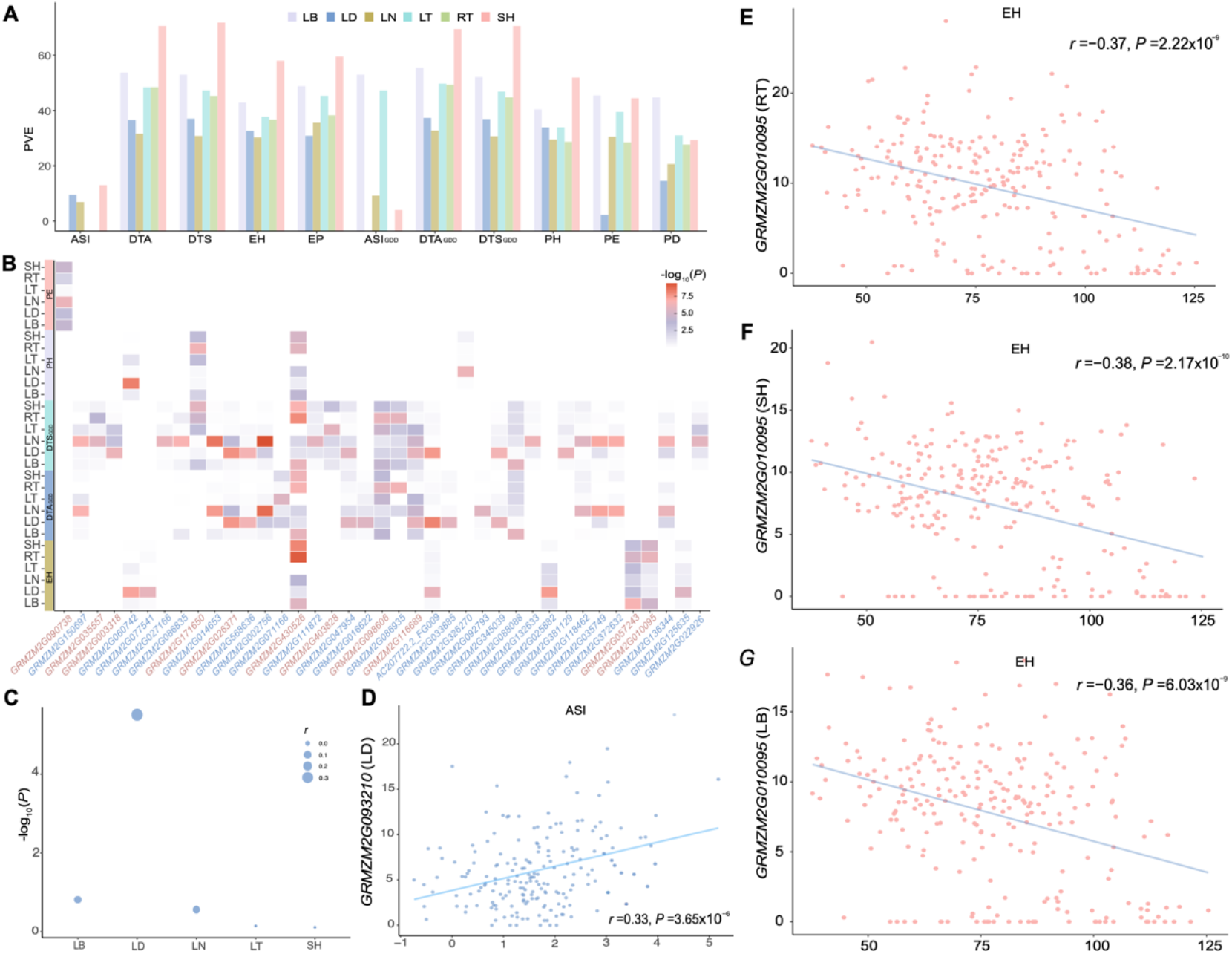
Contribution of tissue-specific gene expression to variation of agronomic trait. (**A**) The proportion of phenotypic variance explained (PVE) by gene expression in seven tissues (**B**) A heatmap illustrating the *P* values of TWAS. Each cell represents a *P* value testing the association between the expression of a gene (x-axis) in specific tissue and a trait (y-axis). The red color highlights genes from category II, while blue represents genes belongs to category I. (**C**) and (**D**) Expression-to-trait correlation and *P*-value testing the association between *GRMZM2G093210* expression in five tissue (RT excluded due to undetectable expression) and ASI variation. (**E**) (**F)** and (**G**) Expression-to-trait correlation maps for *GRMZM2G010095* and EH in RT, SH, and LB.

To explore the relationship between tissue-specific gene expression and variation of agronomic trait, we performed multi-tissue transcriptome-wide association analysis one tissue at a time for the 11 traits. In total, 45 genes were detected with significant association to at least one trait at genome-wide threshold (Fig 4B, Supplementary Table S5). In consistent with previous reports, four out of the 45 genes were reported as important regulators for corresponding trait (Li et al., 2021; Liang et al., 2019). There were drastic differences in the number of associated genes detected for different tissues. For instance, only two genes were identified in RT and LT, while 20 genes were identified in LD. Based on TWAS *P*-values obtained from different tissues (Fig 4B), we classified those genes into two categories. The first category included 31 (69%) genes with expression in only one tissue associated with agronomic trait variation even at a lenient significance threshold correcting for multiple testing of 45 genes (*P*=0.05/45=1.11 × 10^-3^), whereas the second category had 14 (31%) genes with expression in multiple tissues associated with agronomic trait variation. For example, *GRMZM2G093210*, a homologue of *Arabidopsis thaliana* lectin-like receptor kinase 7 essential for pollen development (Wan et al., 2008), showed positive correlation with ASI (TWAS *P* =1.46 × 10^-6^, Fig 4CD) in LD but not the remaining tissues. In contrast, *GRMZM2G010095*, a homologue to *Gossypium hirsutum* C2 NT-type domain-containing protein (Wang et al., 2016), was associated with EH in RT (TWAS *P* = 8.54 × 10^-7^), SH (*P* = 7.51 × 10^-6^) and LB (*P* = 1.64 × 10^-5^, Fig 4E-G). Overall, these results highlighted the role of tissue-specific expression in the biological interpretation of genes associated with agronomic trait with potential benefits in functional characterization of the underlying mechanisms.

### Colocalization of eQTL and QTL associated with agronomic trait

To link QTL associated with agronomic traits and eQTL, we downloaded 2941 previously reported QTL associated with variation of 54 traits, including 44 agronomic traits and ten metabolic traits (Jin et al., 2023; Li et al., 2022; Lipka et al., 2013; Liu et al., 2020; Liu et al., 2021; Owens et al., 2014; Wang et al., 2020) (Materials and Methods, Supplementary Table S6). In total, 1% (33) agronomic trait QTL were colocalized with at least one eQTL, including 21 genes and 15 agronomic traits (Materials and methods, Supplementary Table S7). To evaluate whether the expression of colocalized eGenes was associated with the variation of corresponding agronomic trait, we compared the TWAS *P*-value of eGenes with that from randomly selected genes. We found significantly smaller *P*-values for colocalized eGenes (Supplementary Fig S3), suggesting that they were more likely to be associated with agronomic trait variation. Among these, 11 genes were tissue-specific and associated with variation of 12 agronomic traits, again emphasizing the importance of tissue-specific gene expression in agronomic trait variation. For example, a QTL at chromosome 1: 86,775,549bp was associated with both Zeinoxanthin and *GRMZM2G143202* expression in LD (Fig 5A-C). The G allele increased expression of *GRMZM2G143202* and decreased Zeinoxanthin levels (Fig 5DE).

**Fig 5.**
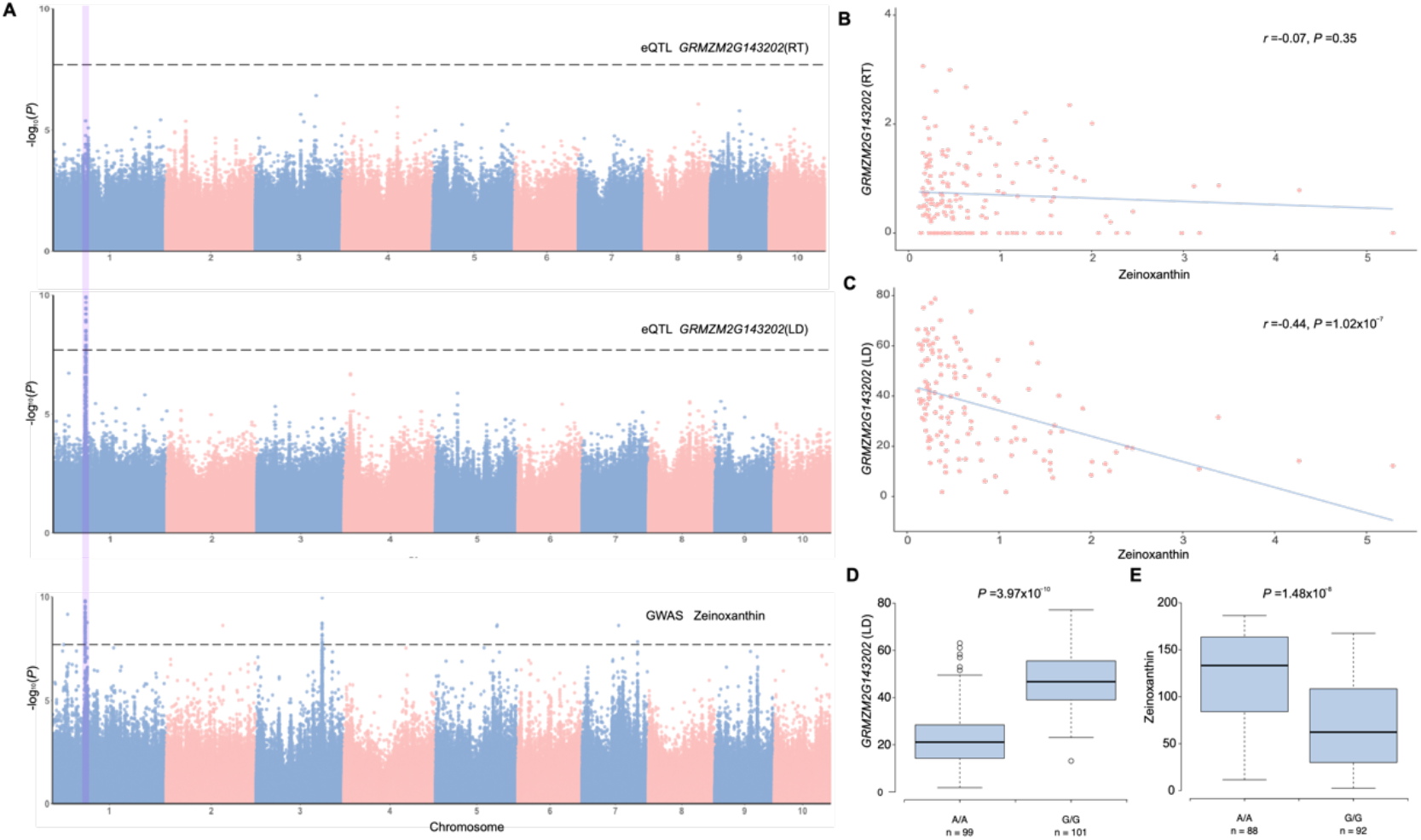
Colocalization of eQTL regulating *GRMZM2G143202* expression in RT and LD, and QTL associated with Zeinoxanthin at chromosome 1: 86,775,549bp. (A) Manhattan plots for *GRMZM2G143202* eQTL mapping in RT, LD, and Zeinoxanthin levels. The vertical line highlights the eQTL and QTL position at chromosome 1: 86,775,549 bp. Gene expression-to-phenotype maps. *GRMZM2G143202* expression in LD **(B)** was negatively correlated with Zeinoxanthin, whereas gene expression in RT **(C)** was not. Genotype-to-phenotype maps. Boxplot of *GRMZM2G143202* expression **(D)** and Zeinoxanthin **(E)** at chromosome 1: 86,775,549bp. *P*-values were obtained from GWAS.

## Discussion

Consistent with previous reports from human GTEx project (Consortium, 2020) and farm animal GTEx project (Liu et al., 2022; Teng et al., 2022), we revealed large variation in the proportion of agronomic trait variance explained by transcriptome differences across tissues (Fig 4A) and detected genes with tissue-specific contribution to agronomic trait (Fig 4B). For instance, SH explained the highest proportion of phenotypic variation for majority of the trait. Through colocalization analysis, we detected eight genes whose expression in SH were associated with a number of agronomic traits. This suggests that SH can serve as a suitable proxy for RNA sequencing when sequencing budget is limited.

One of the motivations to initialize GTEx project was to detect tissue-specific regulatory variants (eQTL) and connected trait-relevant tissues or cell types to complex trait variation. As a pilot study, we detected many tissue-specific genes with reported biological functions to relevant traits (GRMZM2G045318 and GRMZM2G001750 presented above), and found eight candidate genes with potential tissue-specific contribution to agronomic trait variation though literature review (Supplementary Table S8). For example, Rap2.7 encode an AP2 transcription factor, involved in Maize brace roots development and could interact with *ZCN8* (Liang et al., 2019; Li et al., 2019) to regulate flowering time. Our analysis revealed a significant association between the expression of *Rap2.7* in RT and ear height, and a RT-specific eQTL at chromosome 8:131,596,242 bp regulating the expression of *Rap2.7*, suggesting a unique link among the eQTL, *Rap2.7* expression in RT and ear height variation. These findings are valuable for interpreting previous GWAS results and justify a comprehensive functional characterization of the detected genes that have not been connected to variation of agronomic trait. In addition, our analysis revealed 33 genes associated with more than one trait in a tissue specific manner. For example, *AC207722.2_FG009,* annotated as a chlorophyll a-b binding protein, was significantly associated with DTA_GDD_, DTS_GDD_ and, EH in LD (Supplementary Fig S4). Further investigations are warranted to elucidate the underlying mechanisms by which these genes exert their pleiotropic effects and to uncover their broader functional significance in the context of the studied trait.

It is important to note that a limitation of our study is the lack of a comprehensive genotype to phenotype map for maize agronomic traits. Our attempt to assemble such a map was challenged by a lack of publicity in releasing phenotypic data, discordance in genome assembly, and differences in marker density. These limited the power of leveraging multi-tissue gene expression regulatory variant atlas to dissect the genetic basis of agronomic traits in maize. Nevertheless, with an increased amount of public data being available, such as Complete-diallel design plus Unbalanced Breeding-like Inter-Cross (CUBIC) population (Liu et al., 2020) and 1,604 utilized maize inbred lines (Li et al., 2022), the findings present in our study will make a larger contribution to the genetic dissection of maize complex traits. Another potential for the gene expression regulatory variant atlas is enhancing genomic prediction. Integrating tissue-specific genes into extended prediction models in cattle exhibited a modest increase in prediction accuracy compared to traditional models (Fang et al., 2017).

In summary, we present a multi-tissue maize gene expression regulatory variant atlas by analyzing 1771 RNA-seq samples from seven tissues. It represents an extensive reference resource for a detailed characterization of the genetic regulation of gene expression across multiple maize tissues. By establishing links between SNPs, tissue-specific genes expression, and agronomic trait variation, we highlighted a variable contribution from tissue-specific expression to agronomic variation and connected relevant tissues/genes for agronomic traits. The presented results demonstrate the feasibility of uncovering genetic regulatory variants in the maize transcriptome and connecting tissue-specific gene expression to agronomic traits by leveraging public data.

## Material and Method

### Data

298 inbred lines from the maize diversity panel were re-sequenced and called genotypes were downloaded from Kremling et al. (Kremling et al., 2018). 4,260,521 variants with MAF (Minor Allele Frequency) greater than 0.05 were retained for downstream analysis. We downloaded the expression matrix for seven tissues from Kremling et al. (Kremling et al., 2018). Genes expressed (normalized counts> 0) in more than 50% of the individuals were kept for downstream analysis. To compare the effect sizes of detected eQTL across tissues, we standardized the expression values to have mean zero and variance one. BLUPs (Supplementary Table S9) for 30 kernel carotenoid traits, 20 kernel tocochromanol traits and 11 agronomic traits were downloaded from Owens et al. (Owens et al., 2014), Lipka et al. (Lipka et al., 2013), and Peiffer et al. (Peiffer et al., 2014).

### Population structure analysis and expression data clustering analysis

We performed principal component analysis (PCA) with identity-by-state matrix estimated using 4,260,521 SNPs and selected the first two principal components (PCs) to visualize the population structure using the *prcomp* function in R. We visualized sample clusters based on 15,505 genes detected in seven tissues using UMAP approaches in the R package uwot (Melville et al., 2020).

### Detecting tissue-specific gene, function enrichment and evolutionary analysis

We followed the approach presented in Finucane et al (Finucane et al., 2018) to infer tissue-specific expressed genes. First, we scaled the log2-transformed expression (i.e., log2count) of genes to have a mean of zero and variance of one. Then, we fitted the following model to calculate a *t*-statistic for each gene one tissue at a time.

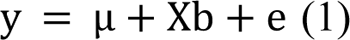

where y is an n × 7 vector of the scaled log2count with n being the sample size and 7 being the number of tissues. μ is the intercept, X is a dummy variable for tissue, where samples from the tested tissue (e.g., RT) were denoted as ‘1’, and samples from remaining tissues (e.g., SH, LB, LT, LD, and LN) were denoted as ‘−1’, b is the corresponding tissue effect, and e is the residual effect. We ranked genes in each tissue according to their *t*-statistic and chose the top 5% as candidate genes. Meanwhile, we filtered these genes by requiring a median fold change of 2 in relation to all the other tissues.

The ClusterProfiler R package (Yu et al., 2012) was used to perform functional enrichment, GO and KEGG, analysis. We obtained the genome of teosinte TIL01 (Zea mays subsp. Parviglumis) from maizeGDB (https://download.maizegdb.org/Zv-TIL01-REFERENCE-PanAnd-1.0/). The nonsynonymous (*d_N_*) and synonymous (*d_S_*) substitutions were computed for all orthologous genes between two entire genomes using the orthologr package (Drost et al., 2015).

### Genome-wide association analysis

GWAS was performed using the *mmscore* function from R package GenABEL (Aulchenko et al., 2007). We performed a conditional analysis where all top-associated SNPs (the SNPs with the highest *P*-value from each association QTL from the initial GWA scan) were included as covariates for additional association signals. We performed this conditional analysis repeatedly until there were no more SNPs that surpassed the significance threshold. This conditional analysis was performed using the *cojo* module (Yang et al., 2012) from GCTA (Yang et al., 2011).

In this study, Bonferroni correction was used to derived the significance threshold. To avoid being over conservative, we estimated the effective number of independent markers (Me) using the GEC software (Li et al., 2012) and derived a less conservative genome-wide significance threshold following 0.05/Me (2.02 × 10^-8^ equivalent to −Log_10_ *P* = 7.69).

### Estimating the contribution from transcriptome variation to agronomic trait variation and transcriptome-wide association analysis

A linear mixed model was used to estimate the amount of phenotypic variance explained by transcriptome variation.

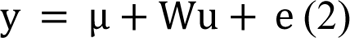

where y is an n × 1 vector of phenotype values with n being the sample size, u is an m × 1 vector of the joint effects of all genes on the phenotype, and e is an n × 1 vector of residuals. W is the corresponding design matrix obtained from a Cholesky decomposition of the transcriptomic relationship matrix (ORM) A, estimated from the transcriptome with OSCA (Zhang et al., 2019). The W satisfied *A* = *WW*^T^; therefore, u and e is random effect with u∼N (0, Iσ^2^_u_) and e∼N (0, Iσ^2^_e_). The proportion of variance explained by transcriptome variation was the intraclass correlation ρ^2^ = σ^2^_u_/(σ^2^_u_ + σ^2^_e_). Transcriptome-wide association analysis (TWAS) was conducted by 6; mixed-linear model approaches (MOA) using OSCA (Zhang et al., 2019). Bonferroni-corrected thresholds (0.05/genes number, 2.58 × 10^-6^) and (0.05/associated genes number, 1.11 × 10^-3^) were used as the significance threshold and lenient significance threshold, respectively.

### Colocalization of eQTL and QTL

We downloaded previously reported maize QTLs from a number of published studies. (Jin et al., 2023; Li et al., 2022; Lipka et al., 2013; Liu et al., 2020; Liu et al., 2021; Owens et al., 2014; Wang et al., 2020). A list of these QTL is summarized in Supplementary Table S10. To adjust differences in reference genome, we lifted over QTL with reported position from maize genome RefGen v4 to AGP v3 using online tools from EnsemblPlants (https://plants.ensembl.org/Zea_mays/Tools/AssemblyConverter?db=core, Supplementary Table S6).

To compile a list of eQTL associated with multi-tissue gene expression, we grouped eQTL from multiple tissue with LD > 0.2 and physical distance < 500kb as a single eQTL. Since LD structure varies between populations in which the trait QTL were detected, trait QTL and eQTL were regarded as colocalized when their physical distance was less than 100 bp.

## Supporting information

Maize-GTEx-SupplementalTable

Maize-GTEx-SupplementaryFig

## Funding

This research was funded by the National Science Foundation of China (32200503), Agricultural Science and Technology Innovation Program (ASTIP-TRIC01) from Chinese Academy Agriculture Sciences, Taishan Young Scholar Program in Shandong Province.

## Author Contributions

Y.Z. and G.Y. designed and supervised this study. M.L., H.S., Z.M., Y.H and L.W. collected the data and implemented the analysis. Y.D. and Y.J. contributed to the statistical analysis. Z.L., F.H, J.Z and R.H. offered valuable input during the design and implementation of this study. Y.Z., S.C. and L.C. wrote the manuscript. All authors have read and approved the manuscript and have no conflicts of interest to declare.

## Acknowledgements

We thank Lingzhao Fang for his constructive comments to improve the manuscript.

## Data availability

The genotypes were downloaded from https://datacommons.cyverse.org/browse/iplant/home/shared/commons_repo/curated/Qi_Sun_Zea_mays_haplotype_map_2018/hmp321_unimputed. The processed expression counts were downloaded from Cyverse Discovery Environment (de.cyverse.org/de/) under the directory: /iplant/home/shared/panzea/dataFromPubs/. Kernel carotenoid BLUPs were download from https://www.ncbi.nlm.nih.gov/pmc/articles/PMC4256781/bin/supp_114.169979_Tabl eS1.xlsx. Kernel tocochromanol trais BLUPs were from https://www.ncbi.nlm.nih.gov/pmc/articles/PMC3737168/bin/supp_g3.113.006148_T ableS1.pdf. The agronomic trait BLUPs were download from http://de.iplantcollaborative.org/dl/d/D94A2327-E2EF-4D44-A7B2-9FC688A2667E/Peiffer2014Genetics_blupPhenos20150325.xlsx. All data from this manuscript is publicly available in previous publications refereed in the method sections.

## Notes

### Competing Interest Statement

The authors have declared no competing interest.

