## Supplementary material for "Comprehensive analyses of 1771 transcriptome from seven tissues enhance genetic and biological interpretations of maize complex traits": Maize-GTEx-SupplementaryFig

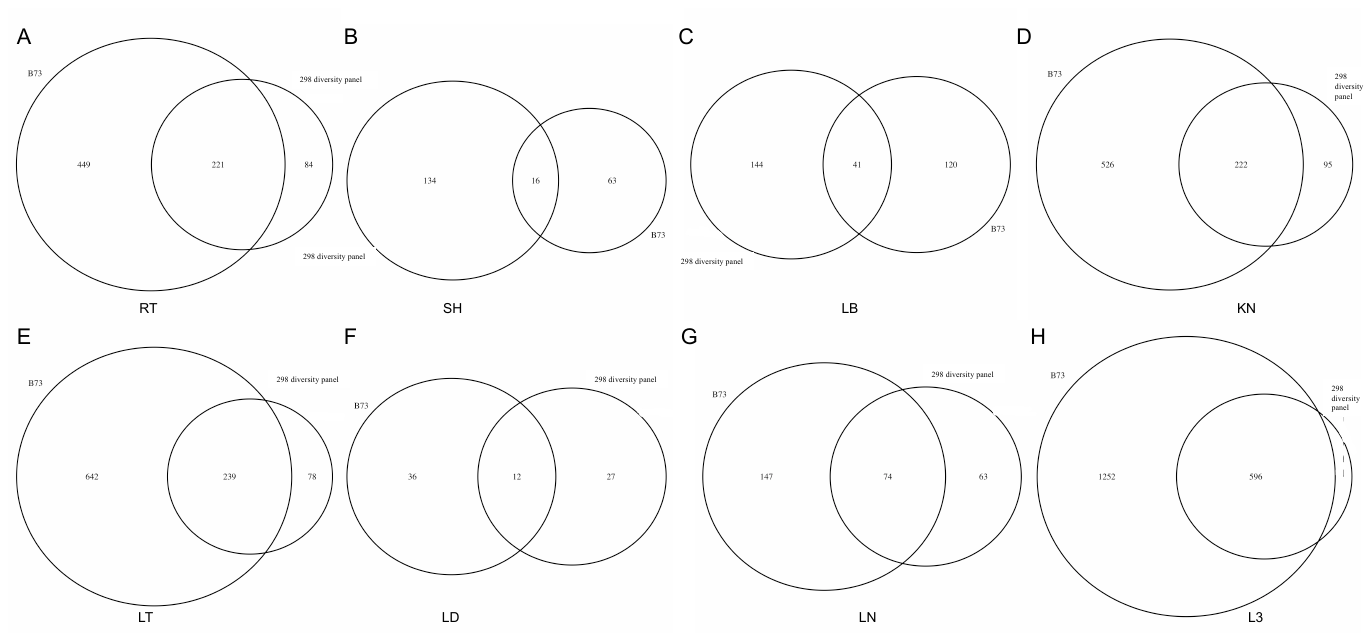

**Fig S1. Venn diagram illustrating overlap of tissue-specific genes from B73 and the maize diversity panel across seven tissues.** L3 stands for LT, LN, and LD

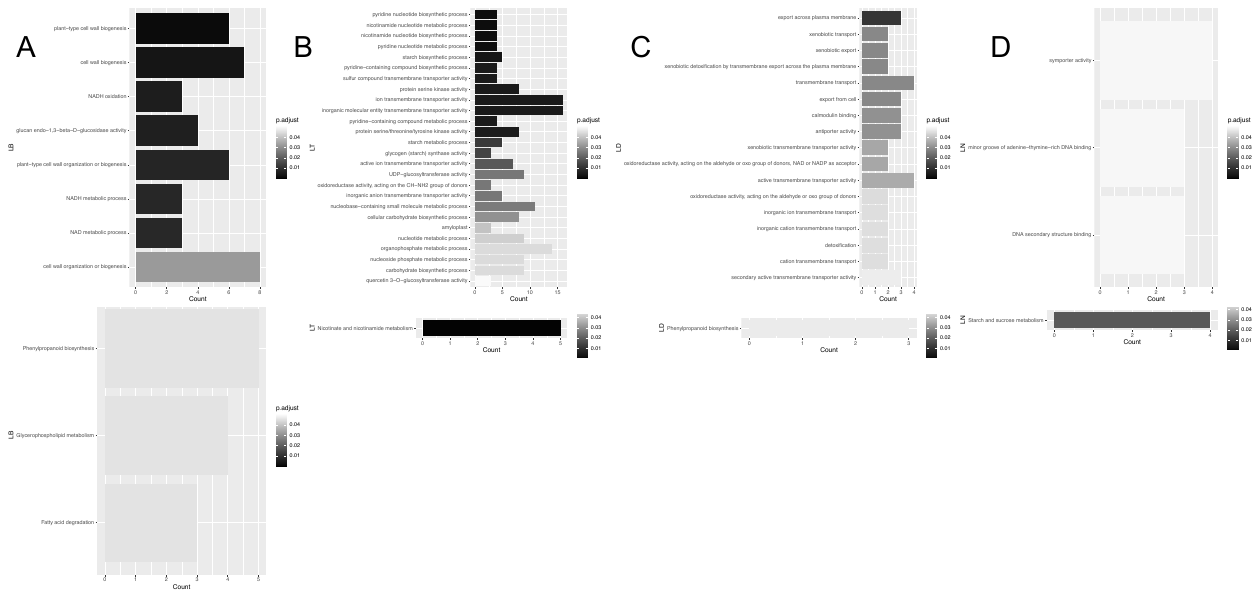

**Fig S2. Gene Ontology (GO) and Kyoto Encyclopedia of Genes and Genomes (KEGG) enrichment analysis of tissue specific genes from LB(A), LT(B), LD(C), and LN(D).** The color corresponds to the enrichment degree (-log10 (adjusted *P*-value)).

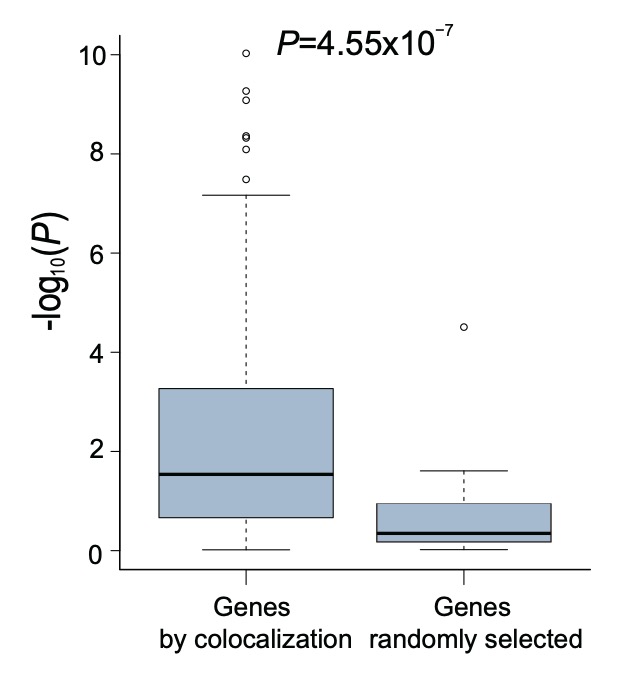

**Fig S3. Comparison of TWAS *P*-values for genes whose eQTL was colocalized with agronomic traits QTLs and randomly selected genes.** *P*-values were calculated using Student’s *t*-test.

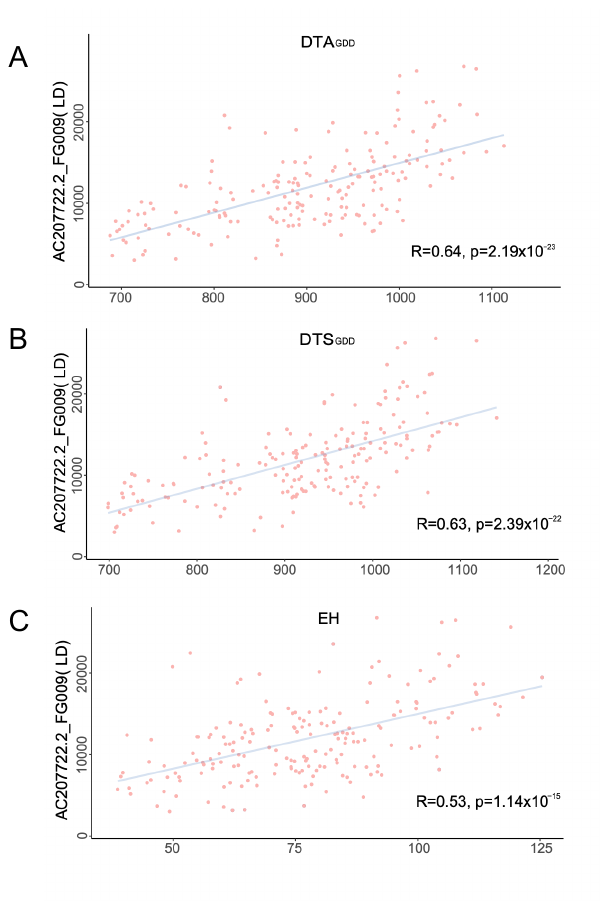

**Fig S4.** **Scatter plot illustration the relationship between *AC207722.2_FG009 expression and*, DTA_GDD_(A), DTS_GDD_(B) and, EH(C) in LD.**
